# Galectin-3 as a new negative checkpoint of the immune response is the key target for effective immunotherapy against prostate cancer

**DOI:** 10.1101/763409

**Authors:** Carolina Tiraboschi, Lucas Gentilini, Felipe M. Jaworski, Enrique Corapi, Carla Velazquez, Anne Chauchereau, Diego J. Laderach, Daniel Compagno

## Abstract

Prostate cancer (PCa) is a major health problem worldwide. Taxol derivatives–based chemotherapies or immunotherapies are usually proposed depending on the symptomatic status. In the case of immunotherapy, tumors develop robust immune escape mechanisms that abolish any protective response. However, Docetaxel has been shown to enhance the effectiveness of immunotherapy in a variety of cancers, but to date, the mechanism is still unknown. Herein, we showed first that Galectin-3 (Gal-3) expressed by prostate tumor cells is the principal immunological checkpoint responsible of the failure of immunotherapy; and that Docetaxel leads to the inhibition of Gal-3 expression in PCa cells as well as in clinical samples of mCRPC patients promoting a Th1 response. We thus optimized a prostate cancer animal model that undergoes surgical resection of the tumor like prostatectomy to mimic what is usually performed in patients. More importantly, using low and nontoxic doses of taxane prior to immunotherapy, we were able to directly impact the activation and proliferation of CD8+ cytotoxic T cells through reducing the number of CD8+CD122+CD28-T cells and highly control tumor recurrence. Thus, Gal-3 expression by PCa cells is a key inhibitor for the success of immunotherapy, and low doses of Docetaxel with noncytotoxic effect on leukocyte survival should be used prior to vaccination for all PCa patients. This combined treatment sequence right after surgery would promote the preconditioning of the tumor microenvironment, allowing for effective anti-tumor immunotherapy and can be transferred rapidly to clinical therapeutic protocols.

## Introduction

Prostate cancer (PCa) is a major cause of suffering and death worldwide (IARC, WHO) (*1*). Early diagnosis and rapid treatment play critical roles in the final outcome. While initial phases with localized and castration-sensitive PCa are curable, those with metastatic and castration-resistant PCa (mCRPC) are not. At this stage, the primary treatment option for symptomatic patients is chemotherapy with Taxol-derived molecules such as Docetaxel. However, 50% of patients develop chemotherapy resistance, and few other therapeutics are available (*2*). It is therefore essential to evaluate alternative approaches to prevent tumor spreading and progression to advanced stages of this disease. In this scenario, immunotherapy, in which the patient’s immune system is targeted to induce an antitumor response, represents an interesting treatment option (*3*).

Immunotherapy is an attractive therapeutic strategy for PCa since tumor cells are not ignored by the immune system, as evidenced by the presence of lymphocyte infiltration in prostate tumors (*4*). These infiltrates are characterized by high levels of regulatory T cells (TReg) (*5–7*). Recent clinical data provide clear evidence of the possibility that antigenic determinants expressed in various types of human tumors could be targeted by autologous T cells and that the optimization of such reactivity could lead to cancer regression (*8–10*). Sipuleucel-T, the first FDA-approved antigen-specific immunotherapy for cancer treatment, is a personalized vaccine based on autologous dendritic cells (DC) that are supposed to activate PAP-specific CD4+ and CD8+ T cells in treated PCa patients (*11*). In fact, Sipuleucel-T is only used for asymptomatic mCRPC patients and induces a 4.1-month improvement in median survival. Furthermore, analysis of the 3-year survival rate demonstrated an 8.7% improvement in patients treated with Sipuleucel-T compared to the placebo group but without effective control of disease progression (*12*). In contrast, GVAX, an allogenic PCa tumor vaccine, failed to demonstrate overall efficiency when compared to Docetaxel therapy. Altogether, the low efficiency of immunotherapies (*13*) demonstrates that prostate tumor cells create a particular microenvironment to evade immune attacks. In this respect, encouraging results have been obtained in clinical trials based on overriding T cell tolerance (*14–18*).

During the last decade, the scientific community demonstrated the involvement of protein-glycan interactions in shaping a tumor-associated immune-suppressive microenvironment (*19*) through multiple mechanisms (*20–27*). While these functions of glycans seem unequivocally described and proven in several experimental settings, the recognition of the glycophenotype by lectins, in particular galectins (Gals), is likely an essential means of tumor-immune tolerance. Interestingly, Gals have been implicated in several situations of immune regulation, with major roles in shaping T cell function in different experimental settings and promoting tumor immune tolerance (*28*). In particular, much attention has been focused on galectin-1 (Gal-1), a member of this family with higher expression levels in PCa and the only galectin whose expression is upregulated during disease progression. Gal-1 seems to have a major effect on neovascularization in PCa (*29*). In contrast, the downregulation of full-length Gal-3 observed in patients apparently matches neither with the definition of Gal-3 as a marker of PCa tumor cell aggressiveness nor with poor marker prognosis for PCa patients (*30–32*). However, because Gal-3 controls the functions of a variety of antitumor immune cells (*33–38*), we decided to further investigate its role in antitumor immune responses. With the goal to transfer our results to clinics, we paid special attention to conditions where a chemotherapy treatment is associated with vaccination since results from clinical trials have shown that Docetaxel-based chemotherapy could promote the effectiveness of immunotherapy in a variety of cancers (*39–44*) as well as in PCa patients (*45–46*). However, to date, neither the mechanism nor any factor has been identified as responsible for this synergic effect of the combinatory therapy protocol. Altogether, these clinical results reveal that much remains to be understood in improving the efficiency of immunotherapy in PCa.

Herein, our results highlight that PCa cell lines at metastatic stage of the disease recover high expression of Gal-3, and under Docetaxel treatment downregulate this new negative checkpoint of the immune responses. More importantly, we demonstrated that the treatment of PCa cell lines or mice-bearing tumors with low and nontoxic doses of Docetaxel (LDD) right after primary tumor resection and prior to immunotherapy promotes the effectiveness of an anti-PCa vaccine through the downregulation of tumor-expressing Gal-3. Such a strategy allows the activation and expansion of antitumor CD8+ cytotoxic T cells to effectively control tumor recurrence, and could be rapidly transfer to clinical protocols for all PCa patients.

## RESULTS

### Negative regulation of Galectin-3 in PCa cell lines delays tumor growth and metastasis development in immunocompetent mice, but not in athymic nude mice

To understand the apparent contradiction between the negative expression of Gal-3 in PCa primary tumors at advanced stages of the disease with the demonstrated roles of this galectin in the development of metastasis and aggressiveness of PCa cells, we designed a murine model using TRAMP-C1 (TC1) with a controlled Gal-3 expression. TC1 cells were generated from transgenic mice with a C57BL/6 genetic background through the insertion of the SV40T antigen under the control of a prostate-specific promoter (TRAMP-C) (*47–48*). These cells can be injected into C57BL/6 mice and develop ectopic prostate tumors. TRAMP-C mice and TC1 cells are widely used murine prostate cancer models that allow the use of syngeneic transplants to study immune responses in immunocompetent animals. We had already standardized a model based on the subcutaneous injection of TC1 to enable us to study various functional aspects of immune cells in PCa (*49–50*). This preclinical model has the advantage that tumor cells and hosts share the same genetic background, which allows the use of immunocompetent mice which is more representative to what generally occurs in clinics. In addition, we generated TC1 expressing different levels of Gal-3 using a lentivirus-derived shRNA expression (Table S1, Fig.1a). On the one hand, control-shRNA transduced cells express high levels of this lectin (TC1-shCtrl or C). On the other hand, cells with low levels of Gal-3 were generated from wild-type cells via transduction with a lentiviral construct encoding a Gal-3–specific shRNA (TC1-shGal-3 or G3). TC1-shGal-3 cells showed a stable 95% decrease of the Gal-3 expression compared with the control (Fig.1a).

**Figure 1:**
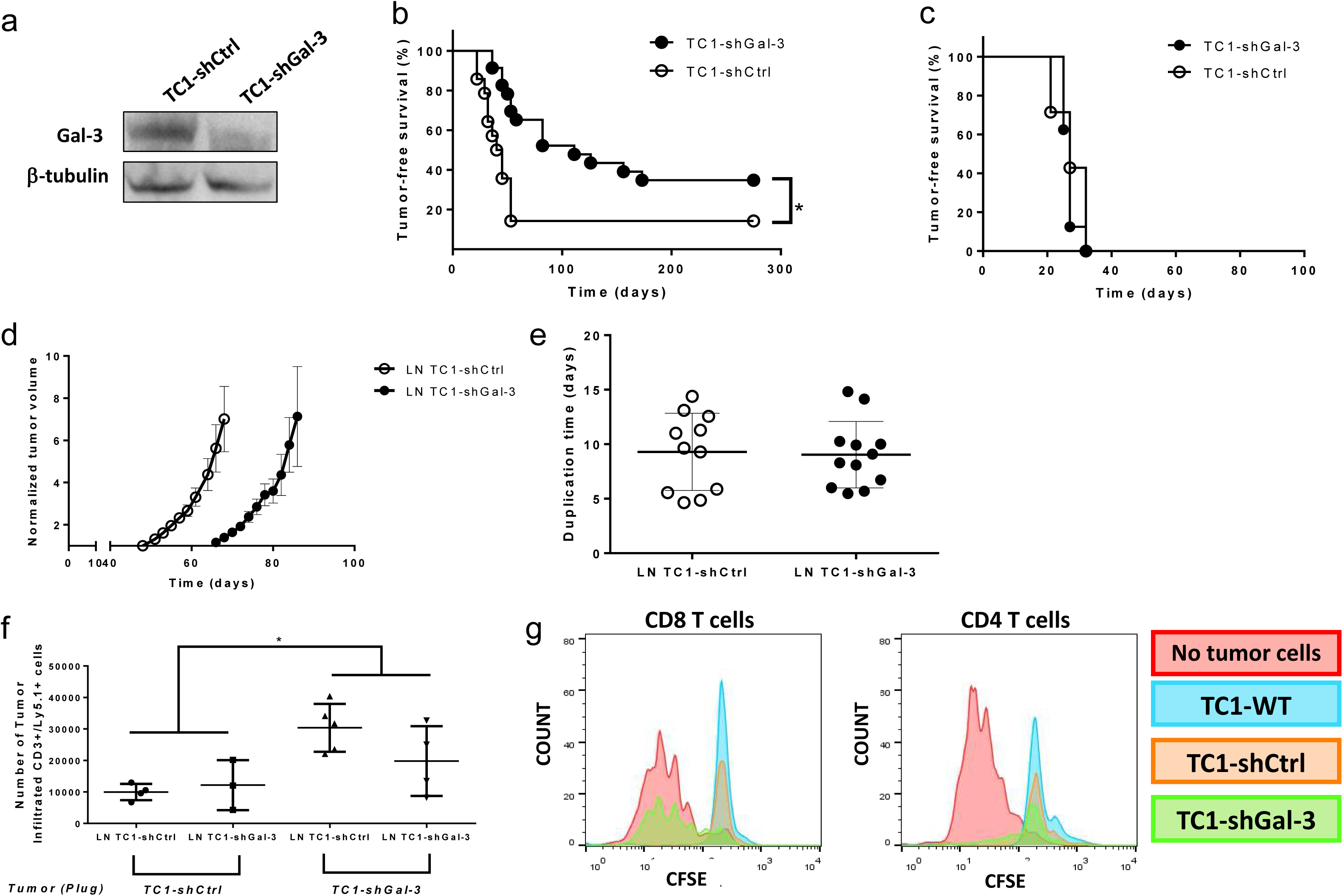
Effect of stable tumor Gal-3 silencing in prostate tumor growth and in tumor cell infiltration. Determination of the level of expression of Gal-3 protein by Western blot (a). Tumor-free survival of TC1-shRNA subcutaneous inoculation in normal (b) or athymic nude (c) C57BL/6 mice. Tumor expressing normal level of Gal-3 (TC1-shCtrl) or silenced Gal-3 (TC1-shGal-3) were adoptive transferred with TC1-shCtrl pre-conditioned lymph node cells (LN TC1-shCtrl) or with TC1-shGal-3 pre-conditioned lymph node cells (LN TC1-shGal-3), (n=12 for each condition) (d-f). Two strains of C57BL/6 mice that differed in the gene variant of the CD45 molecule were used. The Ly5.1 donor strain expresses the CD45.1 variant while the Ly5.2 host strain expresses the CD45.2 variant, which allows the differential analysis of the donor and host cells. Tumor growth (TC1-shCtrl) analysis in Ly5.2 C57BL/6 mice by caliper-measured tumor volume (d). Evaluation of the resulting tumor growth by the time needed to duplicate the tumor volume (d). Analysis by flow cytometry of CD3+ tumor infiltrated cells on day 5 post-adoptive transfer (n=5)(f). *: p<0.05 (Student’s t-test). Proliferation assays were performed with lymph node cells isolated from naïve or immunized mice and co-cultured with autologous adherent spleen cells in presence or not of tumor cells expressing different levels of Gal-3 (Lymphocytes:Tumor cell ratio, 20:1, n=3). Non-immunized mice-lymphocytes were assayed for proliferation after polyclonal in vitro stimulation with coated anti-CD3 antibody (1 μg/ml) for 72 hours, and the proliferation rate was evaluated by dilution of CFSE intensity in the CD8+ or CD4+T cell populations (g).

**Table 1:** Gal-3-silencing in prostate tumor cells by RNA interference or Low/Nontoxic doses of-Docetaxel promotes vaccines effectiveness. Tumor-bearing mice were vaccinated with BM-DC based vaccine loaded with tumor cells lysate expressing different levels of Gal-3 (High level for TC1-shCtrl; or Low level for TC1-shGal-3 and TC1-shCtrl/DTX) as indicated. The treatments with nontoxic doses of Docetaxel (DTX) correspond to a 1nM dose in culture cells during two weeks before processing into lysates or 0.83 mg/kg (i.p. injection) during two weeks, once a week for treated mice. (* p<0.05).

|  | Lysate used in vaccine | Tumor | N | Tumor incidence (%) | Tumor free-survival (days) |
| --- | --- | --- | --- | --- | --- |
| **No V1** | - | TC1-shGal-3 | 14 | 65* | 81±43* |
| **No V2** | - | TC1-shCtrl | 23 | 92 | 39±11 |
| **No V3** | - | TC1-shCtrl/DTX | 10 | 80 | 49±7 |
| **VP1** | TC1-shCtrl | TC1-shGal-3 | 10 | 70 | 116±29 |
| **VP2** | TC1-shGal-3 | TC1-shGal-3 | 10 | 0 | >275 |
| **VP3** | TC1-shCtrl /DTX | TC1-shGal-3 | 10 | 0 | >275 |
| **VP4** | TC1-shCtrl /DTX | TC1-shCtrl | 10 | 90 | 73±35 |
| **VP5** | TC1-shGal-3 | TC1-shCtrl/DTX | 10 | 0 | >275 |
| **VP5-2** | TC1-shGal-3 | TC1-shCtrl /DTX | 5 | 0 | >500 |

We first verify that the absence of Gal-3 expressed by TC1 cells led to the same and already well characterized roles of this galectin namely a decrease in both the tumorigenesis and metastases. Thus, evaluating the tumor growth for several weeks after subcutaneous inoculation of TC1-shGal-3 or TC1-shCtrl in wild-type C57BL/6 mice demonstrated a significant delay of 42±32 days in the tumor apparition and lower tumorigenicity since less animals developed tumors when Gal-3 was silenced in tumor cells (Fig.1b/Table 1 No V1, No V2, *(p<0.01)). Once the tumors had appeared, no difference in tumor duplication time was observed (Table S2). We then analyzed the level of the Gal-3 expression in the resulting tumors to verify that the tumor growth was not due to recovered Gal-3, and showed that Gal-3 was still silenced in TC1-shGal-3-derived tumors compared with the controls (Fig.S1a-b). Also, when we analyzed lately, results show a high decrease in the apparition of metastasis (Table S2). All these results allowed us to validate our TC1 murine model.

**Table 2:** Combination of low and nontoxic doses of Docetaxel and therapeutic immunotherapy leads to the inhibition of prostate tumor recurrence and metastasis. Tumor-bearing mice underwent tumor resection surgery four days before receiving or not i.p. injections of low and nontoxic doses of Docetaxel (DTX: 0.83 mg/kg during two weeks, once a week), and then all tumor-resected mice were vaccinated with BM-DC based vaccine loaded by Gal-3^LOW^-tumor cells lysate (TC1-shGal-3) or not as indicated. Observations were made after the sacrifice of the animals.

|  | Lysate used in vaccine | Tumor | DTX | Primary Tumor recurrence | Metastasis development | Observations |
| --- | --- | --- | --- | --- | --- | --- |
| **No V4** | - | TC1-shCtrl | - | 7/8 | 7/8 | Steatosis |
| **No V5** | - | TC1-shCtrl | + | 4/7 | 1/7 | Splenomegaly + Steatosis |
| **VT1** | TC1-shGal-3 | TC1-shCtrl | - | 4/7 | 4/7 | Increased lymph node size |
| **VT2** | TC1-shGal-3 | TC1-shCtrl | + | 1/7 | 0/7 | Increased lymph node size |

The role of Gal-3 in controlling the function of immune cells in a variety of cancers prompted us to analyze tumor growth in athymic nude mice to further evaluate if the T cell compartment was responsible for these phenotypes. Our results demonstrated neither a delay nor a reduction of tumorigenicity or metastasis development between both tumor conditions (Fig.1c, Table S2). Moreover, Gal-3-silenced tumor growth was faster in nude mice, duplication times were 11±1 and 7±1 days in TC1-shCtrl- and TC1-shGal-3-derived tumors, respectively (Fig.S1, **(p<0.01)). We thus hypothesized that Gal-3 expressed by tumor cells negatively controlled T cell functions to allow faster PCa growth and consequently metastasis development.

### Gal-3 expressed by tumor cells controls the tumor growth kinetic through its action on lymph node cells, decreasing the number of tumor infiltrated T cells without inducing their apoptosis

To further verify our hypothesis of Gal-3 controlling the immune cell functions, we decided to use pre-conditioned lymph node cells to evaluate if the delay on tumor growth could be also obtained by transferring the immunity induced by TC1-shGal-3 (Fig. S2). Briefly, TC1-shCtrl or TC1-shGal-3 cells were subcutaneously injected in C57BL/6 immunocompetent Ly5.1 donor mice in order to pre-stimulate immune cells by tumor cells expressing different levels of Gal-3. After 5 days, total donor lymph node cells (LN TC1-shCtrl or LN TC1-shGal-3) were harvested and adoptive transferred into sub-lethal irradiated Ly5.2 host mice injected the day before with tumor cells expressing wild type level of Gal-3 (TC1-shCtrl). Results confirmed that the absence of Gal-3 expressed by the tumor in the pre-stimulation of donor T cells (LN TC1-shGal-3) promotes a delay of tumor growth in the host mice (Fig.1d), without affecting significantly tumor duplication time (Fig.1e). These results strongly suggested that Gal-3 expressed by the tumor interferes with the establishment of protective immune response.

We thus decided to further study the action of Gal-3 on the tumor infiltrated lymphocytes (TIL) to evaluate whether different priming of T cells has any influence on the number of TIL able to infiltrate tumors expressing wild type or down-regulated Gal-3. For this and as previously described (Fig.S2), we used a adoptive transfer of Ly5.1 donor mice pre-conditioned by TC1-shCtrl or -shGal-3, but this time these lymph node cells were transfered into Ly5.2 mice hosts -bearing TC1-shCtrl or -shGal-3 tumor cells plug. Matrigel-plugs allow us to harvested tumor cells on the day 6 after adoptive transfer to characterize the tumor infiltrated cells.

Results show that the number of CD3+/Ly5.1+ donor TIL were significantly increased in plugs containing Gal-3-silenced tumor (TC1-shGal-3) compared to those containing control tumor cells (TC1-shCtrl)(Fig.1f), suggesting that Gal-3 expressed by the tumor cells is likely to be an inhibitor of TIL infiltration, without any discrimination of how donor lymph node cells were stimulated (LN TC1-shCtrl or LN TC1-shGal-3). Moreover, galectins are also known to induce apoptosis of T cells and this effect could explain this difference in TIL number. We thus assayed for Annexin V/PI labelling and show that Gal-3 expressing tumors (TC1-shCtrl) do not induce major apoptosis of CD3+ cells compared to TC1-shGal-3 tumors (data not shown). Since tumor has been characterized as an immune privilege microenvironment, we hypothesized that the tumor Gal-3 is likely the main inhibitor of T cell proliferation.

### Gal-3 expressed by tumor cells controls the activation and proliferation of activated CD8+ T cells

We then analyzed the effect of the absence of Gal-3 expressed by TC1 cells in promoting the proliferation of immune cells after an in vitro polyclonal activation. Briefly, lymph node cells as the source of T cells were labeled by an intracellular fluorochrome such as CFSE before stimulation in presence of syngeneic splenocytes. After anti-CD3 stimulation, the proliferation of T cells was analyzed by flow cytometry as a 2-time dilution of CFSE. In fact, each pick of CFSE intensity represents each cell division (Fig.1g). As expected, the absence of tumor cells allows for the efficient proliferation of activated T cells, while the presence of both wild-type or control tumor cells (TC1-WT or TC1-shCtrl) inhibits the T cell proliferation of both CD4+ or CD8+T cells. More importantly, the silencing of Gal-3 in tumor cells (TC1-shGal-3) allows the recovery of high proliferation of CD8+T cells but not of CD4+T cells (Fig.1g).

Altogether the results clearly support the hypothesis that Gal-3 expressed by tumor cells is the key factor of the delay of tumor growth by promoting the activation and proliferation of tumor infiltrated CD8+ T cells. More importantly, the expression of this galectins by prostate tumor cells as a new negative checkpoint could control immune cells functions from the begging of the disease to promote an immune ignorance promoting then tumor growth and metastasis.

### Highly effective antitumor vaccine based on Gal-3^LOW^–prostate cancer cell lysate-loaded dendritic cells

To date, immunotherapy has garnered major interest in prostate cancer therapy, but all immunotherapies as Sipuleucel-T (the unique vaccine authorized by the FDA for asymptomatic PCa patients) and other immunotherapies using anti-checkpoint antibodies had failed to show high efficiency against PCa growth or recurrence (*12, 51*). We hypothesized that the expression of Gal-3 by tumor cells could interfere with T cell behavior and thus with vaccine efficiency, and wondered if a therapeutic process similar to Sipuleucel-T using bone marrow–dendritic cells (BM-DC) loaded with a Gal-3^LOW^–PCa cell lysate could be used as an effective vaccine to control PCa tumor growth. To test our hypothesis, we prepared lysates from TC1-shGal-3 or TC1-shCtrl cells by three successive cycles of freezing and thawing to allow for the complete tumor cell lyses. BM-DC were loaded with these lysates independently and matured overnight with adjuvants prior to be used as a vaccine and prior to the inoculation of TC1-shGal-3 cells in C57BL/6 naive mice (Fig.2a).

**Figure 2:**
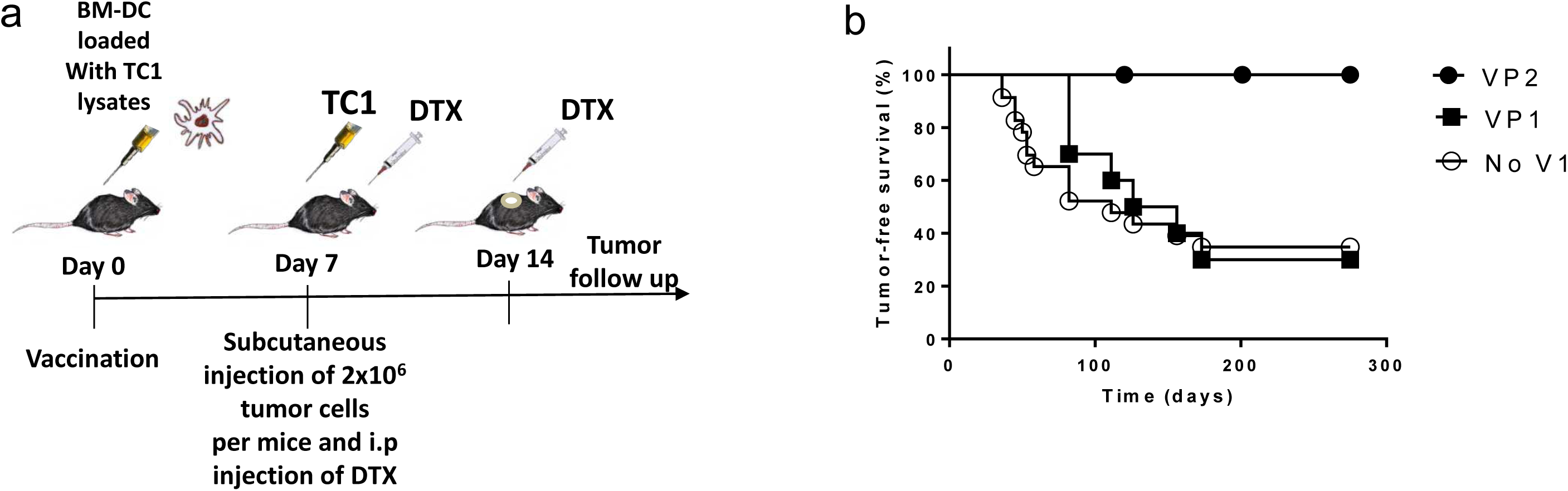
Cellular lysates of Gal-3^LOW^ tumor cells promote an efficient autologous dendritic cell-based vaccine against Gal-3-silenced prostate tumors. Protocol of vaccination uses autologous BM-DC loaded with prostate cancer cell lysates expressing different level of Gal-3 (a). Effect of different cellular lysates on the growth of a Gal-3-silenced TC1 tumor (TC1-shGal-3), (VP1: lysate from TC1-shCtrl; VP2: lysate from TC1-shGal-3; No V1: not vaccinated mice)(b).

Remarkably, the vaccine based on BM-DC loaded with a Gal-3-expressing PCa cell lysate (VP1) allowed a delay in the growth of Gal-3-silenced tumors compared to the unvaccinated mice (No V1), 116±29 days versus 81±43 days respectively (Fig.2b/Table 1: VP1 versus No V1). More importantly, the results also show that the use of Gal-3^LOW^-TC1 cells as a lysate to load BM-DC was sufficient to completely inhibit the tumor growth of cognate Gal-3^LOW^ –TC1 cells, delay superior to 275 days (time of the animal sacrifice) compared to the no vaccination condition (Fig.2b/Table 1: VP2 versus No V1). Altogether, results revealed Gal-3 expressed by PCa cells as a key parameter for the success of immunotherapy.

### Docetaxel treatment promotes the decrease of the Gal-3 expression by prostate tumor cells and in metastatic samples of mCRPC patients

To go further in our study with the goal to translate rapidly research findings into clinical settings, we examined how the expression of Gal-3 could be decreased in patients to promote a pre-conditioning of the tumor microenvironment, allowing the success of an immunotherapy. Docetaxel interferes with microtubule depolymerization, promoting cell cycle arrest and cell death (*52*) and it is widely used as chemotherapeutic agent against PCa in patients. We first analyzed the survival of TC1 cells at different doses of Docetaxel and confirmed that TC1 cells are sensitive to taxane treatments (EC50=8.10±0.03 nM) (Fig.3a). Interestingly, when we analyzed the expression of Gal-1 and Gal-3 (as the most expressed and immunoregulatory galectins in PCa (*29*)), we found that Gal-3 expression strongly decreased in TC1 cells treated with nontoxic doses of Docetaxel compared with cells under vehicle treatment, both in vitro (Fig.3b) and in vivo (Fig.3c). In contrast, the Gal-1 expression was not modified by Docetaxel treatment (Fig.3b-c). More importantly, the Docetaxel-mediated negative regulation of the Gal-3 expression was also confirmed in metastasis samples of mCRPC patients (Fig.3d), and it did not significantly affect the expression of Gal-1. Since Gal-3 is a well-known galectin that could interfere with the immune system (*33–37*) and Docetaxel-based chemotherapy promotes immunotherapy success in PCa patients (*45–46*), we thus hypothesized that Docetaxel acts through Gal-3 silencing in prostate tumor cells, which could be helpful for translational medicine.

**Figure 3:**
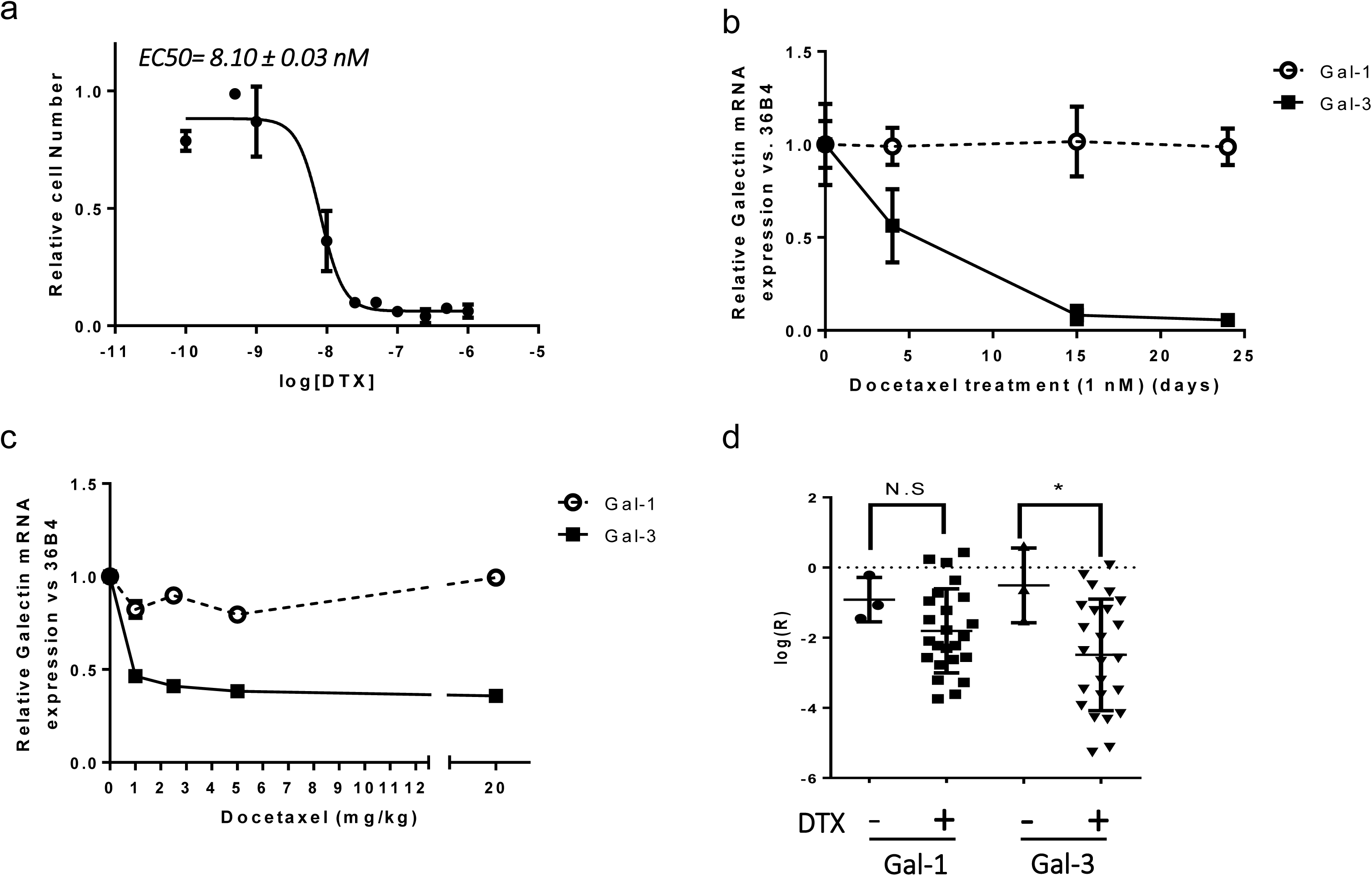
Docetaxel-based chemotherapy induce Gal-3 expression decrease in prostate tumor cells and in mCRPC patients. Dose-dependent cytotoxic effect of Docetaxel in vitro on TC1 (a). Docetaxel promotes Gal-3 silencing in TC1 cell without affecting Gal-1 expression in vitro (b) and in vivo (c), and in mCRPC patient samples using microarray database (GSE35988) (d) (*58*). * p<0.05; N.S: Not significant difference.

### In mCRPC patient samples, Docetaxel-based chemotherapy induces Th1 but not pro-inflammatory gene expression profiles

Although cancer chemotherapy leads to leukocyte aplasia and has always been considered immunosuppressive, numerous clinical and preclinical examples show that certain chemotherapies may increase the efficacy of immunotherapies (*39–46*). Also, the high level of cell death is likely owed to the manner in which chemotherapy promotes a pro-inflammatory microenvironment that achieves additive or synergistic clinical activity with immunotherapy. To date, no study confirms the promotion of an inflammatory microenvironment, especially in PCa patients. Herein, we thus analyzed the expression of a panel of pro-inflammatory genes in metastasis samples of mCRPC patients that have or not undergone chemotherapy protocols. The results in Figure 4 clearly show that the expression of any of the well-characterized pro-inflammatory genes is not modified when analyzed in mCRPC patients treated or untreated with Docetaxel (Fig.4a). To go further in our investigation, we also analyzed cytokines/chemokines gene expression, and confirmed that IL-4, IL-10, TGF-β, and IL-17 genes (as Th2 and Th17 profiles, respectively) showed no significant variation (Fig.4b), while—and more importantly—IL-2 and IFNγ as well as the perforin genes (characteristic of a Th1 profile) significantly increased when mCRPC patients received chemotherapy (Fig.4c). These results strongly suggest that Docetaxel-based chemotherapy could favor the immunotherapy response through inducing Th1 independently to pro-inflammatory genes promotion in mCRPC patients.

**Figure 4:**
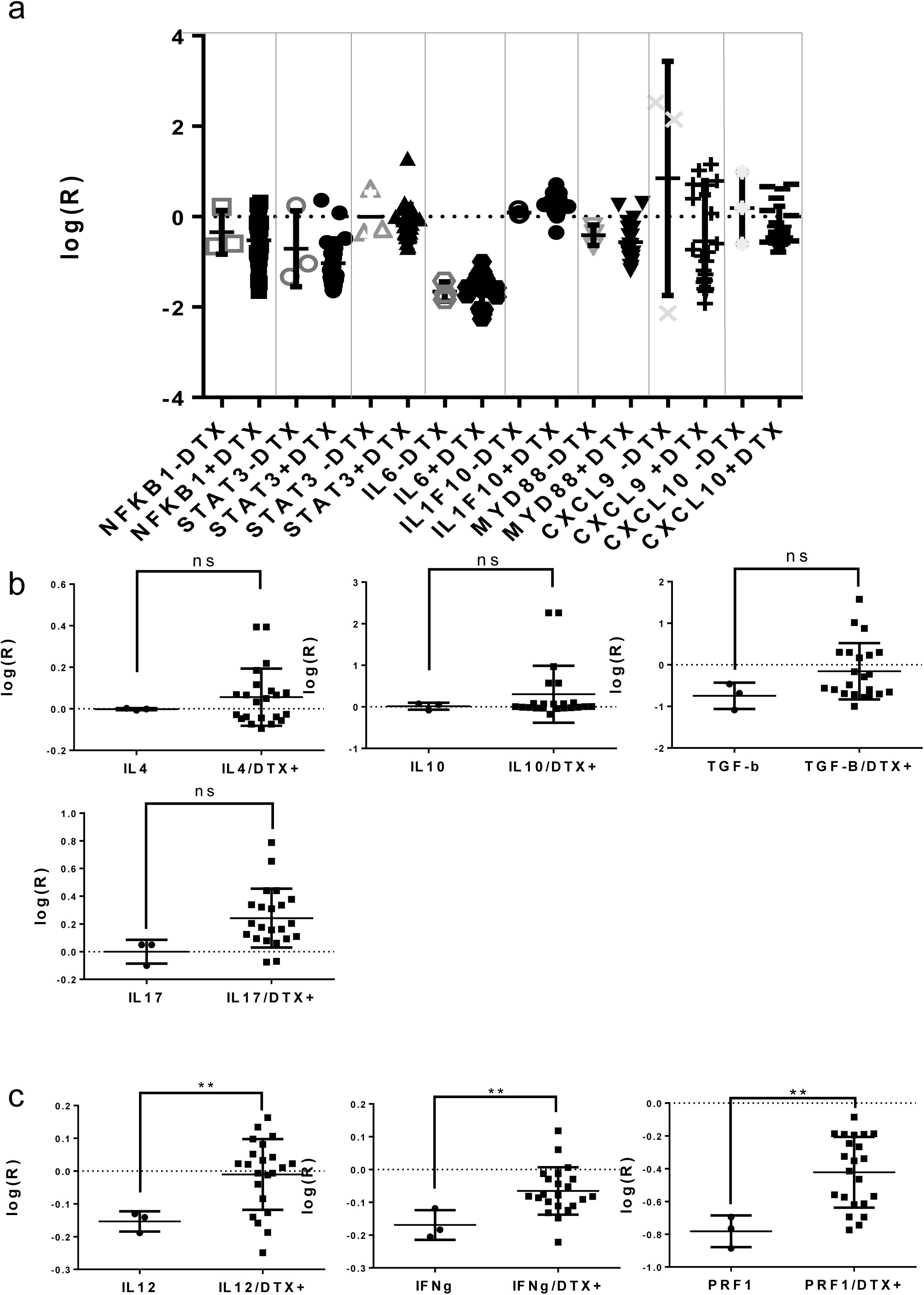
High-throughput analysis of metastasis transcriptome from mCRPC patients, treated or not by Docetaxel-based chemotherapy. Gene expressions in Docetaxel treated (-DTX+) or untreated patients for pro-inflammatory genes (a), for cytokines/chemokines gene expressions for Th2 or Th17 (b) or Th1 profiles (c), using microarray database (GSE35988) (*58*).

### Gal-3 negative regulation in tumor cells is a key factor in the success of immunotherapy

We have shown that a Gal-3-silenced PCa cell lysate used in a BM-DC-based vaccine interferes with the growth of Gal-3^LOW^ tumors using an RNA interference strategy (Fig.2b). With the goal of translational research, we first wondered if low doses of DTX prior to vaccination could strongly decrease the expression of Gal-3 in tumors without affecting the viability of immune cells. We thus sought to analyze the survival immune cells at different doses of DTX. Results in Figure 5 show that in vivo CD8+ T cells (Fig.5a) are sensitive to taxane treatments, while CD4+ T cells are less sensitive (Fig.5b), but no significant effect on the viability of all T cells was observed at doses as low as 0.86 mg/kg. This result prompted us to analyze whether this low dose of Docetaxel-induced Gal-3 decrease enables the same vaccine efficiency seeing with previous tested RNA interference strategy. For this purpose, we tested if a lysate obtained from Docetaxel-pretreated tumor cells (TC1-shCTRL/DTX) could effectively be used in a DC-based vaccine against PCa tumor growth. As shown in Figure 5c, we observed that such a vaccination induced a delay in the growth of Gal-3-expressing tumors but failed to protect the animals (VP4 versus No V3, Fig.5c, Table 1; 73±35 days versus 49±7 days respectively). More importantly, a lysate from Docetaxel-treated TC1 cells (TC1-shCTRL/DTX) did not interfere efficiently with the Gal-3-expressing PCa tumor growth (VP4, Table 1) but rather completely inhibited Gal-3-silenced PCa tumor growth (VP3 and VP5, Fig.5c/Table 1). This inhibition of tumor growth is similar to the anti-tumoral effect obtained with the TC1-shGal-3 vaccine (VP2, Fig.2b/Table 1), and strongly suggests that the main effect of Docetaxel is to interfere with the expression of Gal-3 by tumor cells as well as the RNA interference silencing. Also, a second injection of TC1-shGal-3 cells one year after the VP5 vaccination still showed a complete and longtime protection of mice for tumor growth (Table1). Taken altogether, these results show that the expression of Gal-3 by PCa tumors is a key parameter for the success of immunotherapy since Gal-3 expressed by the tumor is likely the main cause of prostate cancer immunotherapy failure.

**Figure 5:**
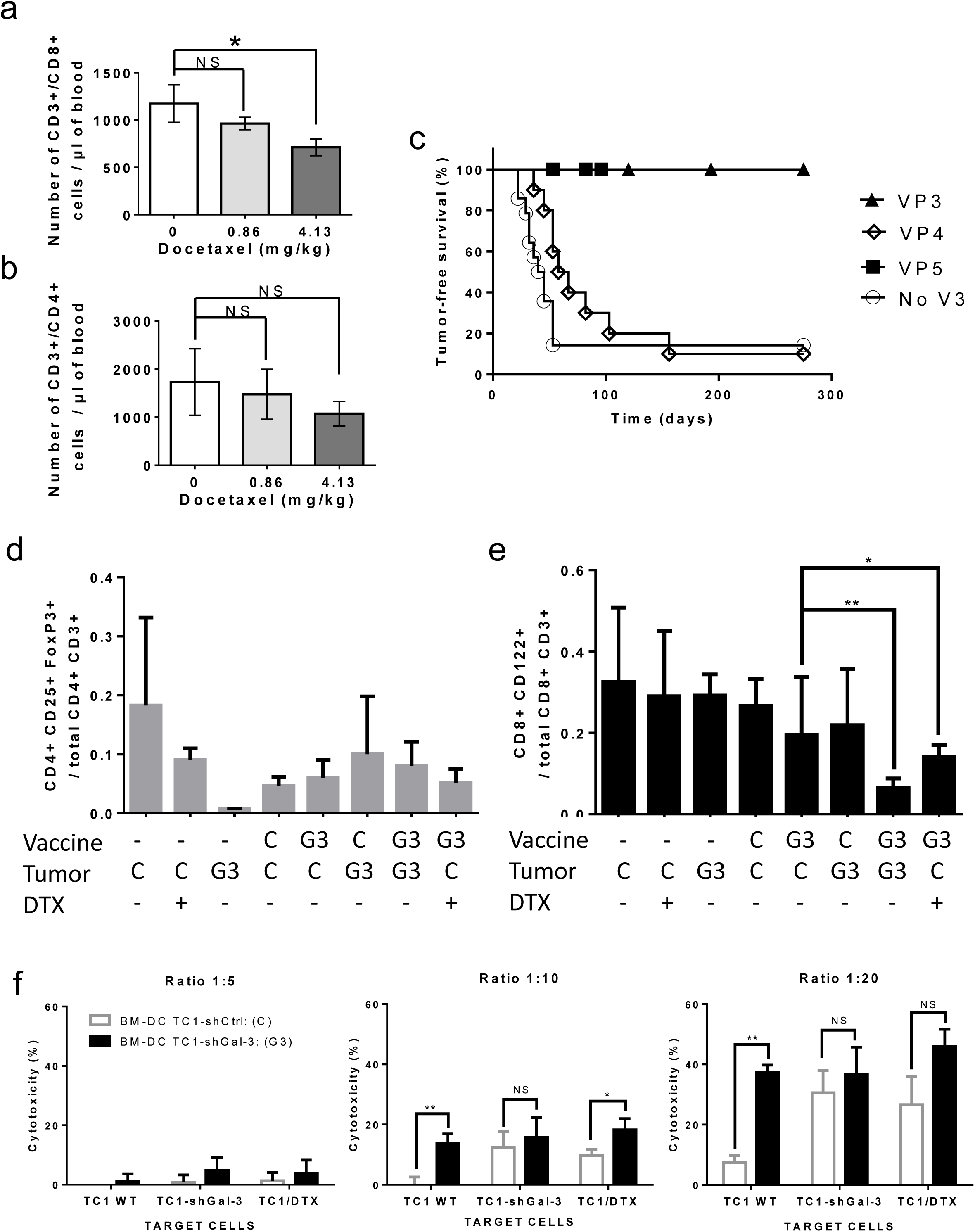
Low and nontoxic doses of Docetaxel induce Gal-3 expression decrease in prostate tumor cells without affecting T cell viability, and promote effective vaccine against prostate tumor. Dose-dependent cytotoxic effect of Docetaxel in vivo on the viability of CD8+ T cells (a) and CD4+ T cells (b). Ex vivo Docetaxel treatment of TC1 cells used as lysate in a DC-based vaccine decreases Gal-3-expressing prostate tumor growth (VP4) and inhibits completely Gal-3-silenced (VP3) or Docetaxel-induced Gal-3 decrease (VP5) prostate tumour growth; No V3: Gal-3-expressing tumor growth on not vaccinated mice (c). Analysis of Regulatory T cells/total corresponding T cells ratio of purified TIL from vaccinated mice bearing different Gal-3-expressing TC1 tumor, on CD4+ (d) and CD8+ (e) compartment. Ex vivo cytotoxic assays of T cells from vaccinated mice against TC1 targets, with control normal (white bars) or negatively-regulated (black bars) expression of Gal-3. Representative histograms and bar graphs of three independent experiments are shown (f).

### Vaccination with Gal-3^LOW^–prostate cancer cell lysate loaded DC activates cytotoxic CD8+T cells and reduces the number of CD8^+^-but not CD4^+^-regulatory T cells

To continue our analysis and verify our hypothesis that the vaccination success was due to the effective activation of an anticancer immune response in absence of Gal-3, we performed tumor infiltration analysis and cytotoxicity assays with T cells from immunized mice as additional functional studies. Since PCa is also characterized by a high level of regulatory T cell (TReg) infiltration (*5–7*), the induction and differentiation of TReg by Gal-3 expressed by tumor cells are other important parameters that may impact tumor development. It is well known that CD4+CD25+Foxp3+ regulatory T cells (CD4TReg) suppress cytotoxic CD8+T cell function. Their inhibitory function depends on Galectin expression (e.g., essentially Gal-1 (*22*)). Since the Gal-1 expression is not modified in TC1-shGal-3 and in Docetaxel-treated patients, we thus wondered whether Gal-3 could have the same function to promote PCa immune tolerance. Then, we decided to further study the levels of CD4TReg (CD4+CD25+Foxp3+) and CD8TReg (CD8+CD122+CD28-) in the prostate tumor microenvironment depending on the level of Gal-3 expression. For this, we vaccinated mice with different conditions of Gal-3-expressing tumors, treated or not with Docetaxel to allow Gal-3 negative regulation, and then evaluated in the immune cells infiltration the TReg/total T cells ratios: CD4TReg versus total CD4+T cells or CD8TReg versus total CD8+T cells (Fig.5d-e). Despite the slight and insignificant differences observed in the CD4+T cell population (Fig.5d), the ratio between CD8TReg versus total CD8+T cells strongly decreased when the PCa tumors lacked Gal-3 (Tumor G3 or DTX-treated tumors (C, DTX +)) and in the mice vaccinated in the absence of Gal-3 (vaccine G3) (Fig.5e). Interestingly, the level of Gal-3 should be negatively controlled in the lysate that loaded BM-DCs (Vaccine G3) to effectively control the tumor growth. In fact, vaccination with the lysate from TC1-shCtrl (vaccine C) only delayed tumor growth and was unable to decrease the ratio of CD8TReg to total CD8+ T cells (VP1 vs No V1, Table 1 16±29 vs 81±43 and Fig.5e respectively), unlike the vaccination with the lysate from both Gal-3-negatively regulated TC1 (VP2 and VP3, Table 1 and Fig.5c). Moreover, the results also confirm that Gal-3 expressed by tumor cells (Tumor C, DTX-) plays a key role in inhibiting CD8+T cell proliferation, allowing high levels of CD8TReg to control antitumor immune responses and thus allow PCa tumor growth. More importantly, the success of the vaccination is likely dependent on the Gal-3 negative status of the tumor (G3 or (C, DTX+)). To continue this functional study, we decided to test if a Gal-3 expression by tumors could affect the resultant cytotoxicity of the activated CD8+T cells. For this, we analyzed antitumor cytotoxicity after vaccination using LDH release assays (Fig.5f). The results show that vaccination with DC loaded with a TC1-shCtrl lysate (white bars) promotes the killing of cognate target cells (e.g., TC1 WT that express a normal level of Gal-3), while vaccination with DC loaded with Gal-3-silenced TC1 (TC1-shGal-3 or TC1/DTX) is much more effective (Fig.5f). Moreover, vaccination with DC loaded by the lysate from TC1 that does and does not express high levels of Gal-3 is capable of inducing the death of Gal-3-silenced TC1 targets (TC1-shGal-3 and TC1/DTX). Altogether, these results support the assumption that the negative regulation of Gal-3 in prostate tumor cells is required to negatively control the number of CD8TReg cells and thus allow high antitumor CD8+T lymphocyte proliferation and cytotoxicity. We thus hypothesized that Gal-3 is a negative checkpoint of the immune response controlling the cytotoxic function of activated T cells, and could be targeted by a preconditioning treatment with low and nontoxic doses of DTX (LDD) prior to vaccination.

### A preconditioning treatment with low and nontoxic doses of Docetaxel prior to immunotherapy is the key to lead to an effective therapeutic vaccine against prostate cancer

We had confirmed that the expression of Gal-3 by the tumor could be responsible for the failure of immunotherapy against PCa. Since LDD interfere with the expression of this galectin by prostate tumor cells but do not promote cell death (neither tumor cells and, more importantly, nor immune cells), we decided to test whether an in vivo treatment with this taxane prior to vaccination could protect prostate tumor–bearing animals by improving the immunotherapy efficiency. For this purpose we used our preclinical model including a tumor resection surgery that mimics prostatectomy protocol and allowing for the evaluation of tumor recurrence. Then, the tumor-resected mice were treated two times and once a week with LDD (0.83 mg/kg, 47-time less compared to the corresponding chemotherapeutic doses used in humans, (http://www.fda.gov/cder/guidance/index.html). This preconditioning treatment on the day 4 after the tumor resection (DR, Figure 6) allows negative Gal-3 regulation in the remaining tumor cells (likely circulating tumor cells) before vaccination with an autologous BM-DC loaded by a Gal-3^LOW^–prostate cancer cell lysate (TC1-shGal-3) (Fig.6a/Table 2). Without any vaccination, the results in Table 2 show first that LDD-induced decrease of the Gal-3 expression in tumor cells is insufficient to control Gal-3-expressing tumor growth, since 7 from 8 treated animals show recurrence of the primary tumor and metastasis development (No V5 versus No V4, Table 2:) but controls metastasis development as well as with Gal-3-silenced tumors (Fig.S1c). Second, vaccination with an autologous BM-DC loaded with a Gal-3^LOW^–tumor cells lysate is inefficient over a long period to inhibit tumor recurrence after the tumor resection (VT1, Table 2). More importantly, these results reveal that these low doses of Docetaxel preconditioning right after primary tumor resection and prior to vaccination is essential to allow immunotherapy to control PCa tumor growth, as demonstrated by the absence of tumor recurrence in a large majority of mice (1/7) treated with the combinatory approach (VT2, Table 2). These results show that Gal-3 expressed by prostate tumor cells influences directly neither the metastatic progression nor the tumorigenicity of PCa cell lines, but more importantly, allows CD8+CD122+CD28-T cell differentiation to inhibit activated antitumor CD8+ T cells (Figure 6b). Thus, Docetaxel-inducing Gal-3 negative regulation is the main factor in the chemotherapy promotion of immunotherapy against PCa. Finally, chemotherapy based on low and nontoxic doses of Docetaxel prior to immunotherapy allows tumor-free outcomes in a large majority of animals. This protocol could be easily transferable to clinical settings to treat all PCa patients as soon as they suffered a prostatectomy surgery.

**Figure 6:**
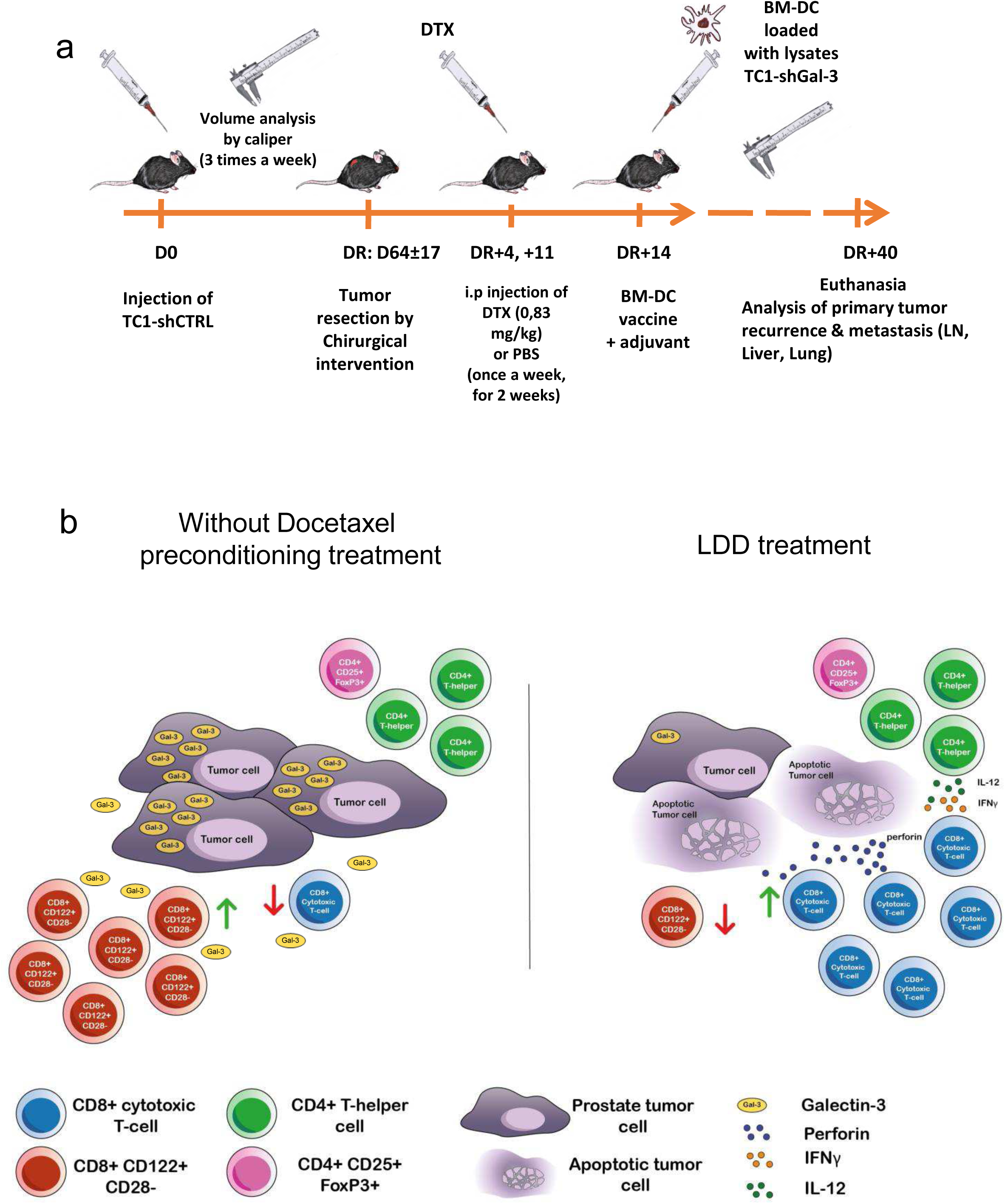
In vivo treatment with low and nontoxic doses of Docetaxel prior to a BM-DC vaccination leads to effective immunotherapy against PCa, preventing the tumor recurrence. Protocol of surgical tumor resection followed by autologous BM-DC vaccination, combined or not with Docetaxel treatment prior to vaccination, and evaluation of tumor recurrence. Schematic model of the effects of Docetaxel preconditioning treatment combined with immunotherapy in vivo (a). A two-week treatment with low and nontoxic doses of Docetaxel (LDD) leads to a strong decrease in the expression of Gal-3 by tumor cells. When vaccinated with BM-DC loaded with Gal-3Low-tumor cell lysate, mice subjected to LDD treatment (right) showed a decrease in the CD8+CD122+CD28-/total CD8+ cells ratio in comparison to non-pre-treated mice (left), while the CD4+CD25+FoxP3+ (CD4Treg)/total CD4+ cells ratio remained constant. Moreover, metastatic samples of mCRPC patients treated with this chemotherapy presented an increased expression of genes that favor an effective cytotoxic response, like perforin and Th1 profile cytokines/chemokines (b).

## DISCUSSION

In this study, we addressed conflicting findings of roles of Gal-3 and its down-regulation in primary prostate cancer samples. We first demonstrated that Gal-3 is responsible for the aggressiveness of prostate cancer cells and the metastasis development through its control of the immune system. Then, we also described how Docetaxel-based chemotherapy positively affects the effectiveness of immunotherapy against PCa. Our results show that Docetaxel treatment negatively regulates the Gal-3 expression on tumor cells. This biological effect results in potentiating the response of the immune system to an effective anti-PCa vaccination, positioning Gal-3 as a major negative checkpoint of the immune response that allows PCa growth and aggressiveness. These results match previous bibliographic data suggesting a correlation between the level of the Gal-3 expression by the tumor and poor prognosis for PCa patients (*30–32*), and allow us to propose a functional combinatory therapeutic protocol against PCa recurrence for all patients.

Galectins have already been shown to be proteins involved in controlling immune responses in a broad range of diseases (*28, 38*). Most reports on cancer have concentrated on Gal-1 and its effect on immune escape, but it must be emphasized that different galectin members can have different and sometimes opposite effects on T cell behavior (*20*). Gal-3 has been shown to be strongly expressed by PCa primary tumors at the beginning of the disease and decreases to the complete turnout of its expression at advanced stages of the disease (*29, 53*), suggesting its main function might be to control the priming of the antitumor immune response. We showed here that this particular galectin recovers its expression in metastasis samples of mCRPC patients, confirming the correlation between Gal-3 expression and poor prognosis for PCa patients. In this study, we showed for the first time that Gal-3 is also required for prostate tumor cells to establish and maintain an immune tolerance and that this occurs through inducing the deregulation of CD8+T cell cytotoxic responses. Ideally, an effective antitumor vaccine requires the correct priming of naive T cells, which, in turn, acquires effector functions that enable the eradication of tumor cells. The data in the literature clearly demonstrate that cytotoxic CD8+T cells are the main cell type whose presence in infiltrates is associated with better prognosis in all types of cancers (*54*). Our results show for the first time that Gal-3 negative regulation in tumor cells (accomplished by two different strategies: Docetaxel treatment or lentivirus-drived stable RNA interference) allows the efficient activation and proliferation of CD8+ cytotoxic T cells by decreasing the ratio between CD8+CD122+CD28-regulatory T cells and total CD8+ T cells. Additionally, it has been reported that Gal-3 in a tumor microenvironment could inhibit CD4 and CD8 T cell functions (*55*). More importantly, Gal-3 secreted by tumor cells could sequester the INFγ in the stroma (*37*), inhibiting the function of this cytokine that is required for the correct polarization of Th1 cells and the cytotoxic activity of T cells. Our high-throughput analysis of gene expression shows that Docetaxel chemotherapy decreases Gal-3 expression in mCRPC patient samples and promotes the expression of Th1 genes and cytoxic genes such as perforin. Altogether, these data support also tumor-derived Gal-3 as a key negative checkpoint of the anti-PCa immune response, promoting the expansion of CD8+CD122+CD28-regulatory T cells that finally inhibits the cytotoxic functions of activated and antitumor CD8+ T cells.

Recently, it has been observed that Taxol derivates–based chemotherapy has a positive influence on cancer immunotherapy. In fact, some reports revealed that Docetaxel treatment promotes the survival of activated T cells in colon (*40*), Lewis lung (*39*), and metastatic breast (*43*) cancers. This prompted us to evaluate the effect of Docetaxel in tumor cells and leukocytes. We found that low and nontoxic doses of this taxane, as a 47-time lower doses that currently used in chemotherapeutic protocols, neither promote lymphopenia nor induce the death of tumor cells (Fig. 5a-b or Fig.3a, respectively). However, treatment with these low doses of Docetaxel strongly decreases the expression of Gal-3 by tumor cells, both in vitro and in vivo, leading to a strong decrease of CD8+ regulatory T cells and thus a high efficiency of antitumor vaccination protocols. This enhanced immune response by a Docetaxel-induced Gal-3 decrease in tumor cells involves an antitumor CD8+T cell expansion with effective cytotoxic functions. This finding can have a great impact on the development of immunotherapies for PCa. Given the limited success of the only immunotherapy approved for mCRPC patients (Sipuleucel-T; overall survival of 4.1 months (*12*)) and the absence of a response of all other immunotherapies against PCa, our results suggest that the Gal-3 expression by tumor cells or circulating prostate tumor cells could be one of the reasons for the ineffectiveness of these strategies for treating PCa patients. Finally, our results suggest that the efficiency of a DC-based vaccine against PCa strongly depends on the Gal-3 status of the tumor. Since primary tumors in advanced PCa are mostly Gal-3 downregulated, it is conceivable that these phases of the disease are favorable to immunotherapy. However, the decreased expression of Gal-3 was only identified in primary prostate tumors but not in mCRPC samples and no data exist on the level of expression of this galectin in corresponding circulating tumor cells (CTC), the remaining cells after a tumor resection or prostatectomy. It is likely that the expression of Gal-3 might be controlled before PCa patients undergo immunotherapy protocols but that the reduced number of CTC in PCa patients does not easily allow this kind of pre-analysis (*56*). Gal-3 could be then used not only as a bad prognosis for PCa patients, but also as a marker of immunotherapy resistance. We thus propose that patients should take advantage of a pretreatment with low and nontoxic doses of Docetaxel as a preconditioning treatment to decrease the Gal-3 expression by the remaining tumor cells and prior to vaccination to improve immunotherapy success.

